# A genome-wide circular RNA transcriptome in Rat

**DOI:** 10.1101/2021.02.20.432122

**Authors:** Disha Sharma, Paras Sehgal, Sridhar Sivasubbu, Vinod Scaria

## Abstract

**Background:** Circular RNAs are a novel class of non-coding RNAs that backsplice from 5’ donor site and 3’ acceptor site to form a circular structure. A number of circRNAs have been discovered in model organisms including human, mouse, Drosophila, among other organisms. There are a few candidate-based studies on circular RNAs in rat, a well studied model organism. The availability of a recent dataset of transcriptomes encompassing 11 tissues, 4 developmental stages and 2 genders motivated us to explore the landscape of circular RNAs in the organism.

**Methodology:** In order to understand the difference among different pipelines, we have used the same bodymap RNA sequencing dataset. A number of pipelines have been published to identify the backsplice junctions for the discovery of circRNAs but studies comparing these tools have suggested that a combination of tools would be a better approach to identify high-confidence circular RNAs. We employed 5 different combinations of tools including tophat_CIRCexplorer2, segemehl_CIRCexplorer2, star_CIRCexplorer, Bowtie2_findcirc and Bowtie2_findcirc (noHisat2) to identify circular RNAs from the dataset.

**Results:** Our analysis identified a number of tissue-specific, developmental stage specific and gender specific circular RNAs. We further independently validated 16 circRNA junctions out of 24 selected candidates in 5 tissue samples. We additionally estimated the quantitative expression of 5 circRNA candidates using real-time PCR and our analysis suggests 3 candidates as tissue-enriched

**Conclusion:** This study is one of the most comprehensive studies that provides a circular RNA transcriptome as well as to understand the difference among different computational pipelines in Rat.

## Introduction

The last decade has witnessed tremendous advances in technology which has enabled an unprecedented opportunity to understand genomes. RNA-sequencing is one of the approaches which have tremendously improved our depth and breadth of understanding transcripts and transcript isoforms at resolutions previously few could fathom. This consequently has helped in the discovery of different classes of RNAs, including small RNAs like miRNAs, siRNAs and long non-coding RNAs including antisense RNAs, intronic RNAs, lncRNAs. It is now well understood how these different classes of RNAs play a key role in the regulation of gene expression and regulate key developmental and pathophysiological processes(Holdt et al., 2018; Qu et al., 2017; Rybak-Wolf et al., 2015).

Circular RNAs, an emerging class of transcript isoforms, which have been rediscovered recently and found to be ubiquitous in all cellular organisms(Salzman, 2016; Zhao et al., 2019). These isoforms are produced by a unique back-splicing of exons and therefore could be classified as a splice isoform. Although earlier considered as splicing errors, it has now been shown that circular RNAs are present in a number of organisms including model organisms like humans(Long et al., 2018; Memczak et al., 2013; Nicolet et al., 2018; L. Song & Xiao, 2018; Z. Zhang et al., 2018), worms(Cortés-López et al., 2018), Drosophila(Cortés-López et al., 2018; Westholm et al., 2014),mouse(Feng et al., 2018; Singh et al., 2018; J. Zhang et al., 2018), chicken(Shen et al., 2019; Y. Wang et al., n.d.) and pig(Guo et al., 2019; Liang et al., 2020; Mester-Tonczar et al., 2020). Recent studies have also suggested that circular RNAs might have a role in regulation of gene expression of protein-coding genes(X. Li et al., 2018). Circular RNAs have also been shown to have multiple miRNA binding sites, suggesting their role as miRNA sponges(Lin & Chen, 2018; X. Li et al., 2018; X. Wang et al., 2018; Z. Wu et al., 2018). Different reports have also suggested the role of circular RNAs in disease and their potential as disease biomarkers(Y. Li et al., 2018; Sheng et al., 2018; Weng et al., 2019). Memczak *et al* have identified 1900 transcripts from the exonic, intronic and intergenic region(Memczak et al., 2013). Jeck et al also identified exonuclease treated transcripts >25,000(Jeck et al., 2013; Memczak et al., 2013). Circular RNA are known to exist as cell-line specific and tissue-specific isoforms(Coscujuela Tarrero et al., 2018; C. Zhou et al., 2017) and are also known to have conserved splice sites.

Till now, several computational methods for the detection of back-splice events from RNA-Seq data have been developed, such as CIRCexplorer(X. Zhang et al., 2019; X.-O. Zhang et al., 2014), testrealign/segemehl(Hoffmann et al., 2014; X.-O. Zhang et al., 2014), circRNA_finder(Westholm et al., 2014), find_circ(Memczak et al., 2013; Westholm et al., 2014), CIRI(Gao et al., 2015), UROBORUS(Gao et al., 2015; X. Song et al., 2016), NCLscan(Chuang et al., 2016), PTESFinder(Izuogu et al., 2016), KNIFE(Izuogu et al., 2016; Szabo et al., 2015), Pcirc_finder(L. Chen et al., 2016) and Acfs(You & Conrad, 2016). Each of the algorithms for circRNA identification rely on different approaches including different read aligners, requirement of genome annotations and also in many cases in the output formats. In a study by Hansen *et al*, 2015(Hansen et al., 2016), five circular RNA identification tools including CIRCexplorer(Hansen et al., 2016; X.-O. Zhang et al., 2014), circRNA_finder(Westholm et al., 2014), CIRI(Gao et al., 2015), find_circ(Memczak et al., 2013) and MapSplice(Memczak et al., 2013; K. Wang et al., 2010) were compared for the levels of false-positives and sensitivity. The comparison suggested that the high number of circRNAs from a pipeline does not necessarily mean true positive candidates and one single method is not reliable. The study also concluded that it is perhaps better to use a combination of two or more methods to increase the robustness of circRNA detection, increase sensitivity and reduce false negative and false-positive predictions. CirComPara is one such pipeline that by default uses four circRNA prediction tools including CIRCexplorer2, CIRI, find_circ and test-realign with different combinations for different aligners(Gaffo et al., 2017).

Studies have shown that humans, mice and rats share high levels of genetic conservation among each other. Rat, the first mammalian species domesticated for research purpose(Kuramoto et al., 2012)is the most widely studied experimental model so far in medical research and an excellent model for cardiovascular disease, stroke(Gaffo et al., 2017; Rubattu et al., 1996) and hypertension(Bier et al., 2018; Redina & Markel, 2018; Sung et al., 2018) and a variety of genetic stocks(Alam et al., 2014, 2015; Serikawa et al., 2015). In many cases rats were preferred to the mouse to model learning, memory and cognitive research(Vorhees & Williams, 2014). Rat is also a primary model for mechanistic studies in the field of human reproduction and a standard model for physiological and toxicological studies (Martin et al., 2011; Vorhees & Williams, 2014).

Yu *et al* have provided an rRNA depleted rat RNA-sequencing transcriptome map for 11 tissues and 4 developmental stages (2 week-old, 6 week-old, 21 week-old and 104 week-old). Subsequent studies on rat circular RNA transcriptome have used this dataset to identify circular RNAs using CIRI(T. Zhou et al., 2018) but the predicted candidates were not independently validated experimentally.

In this study, we used publicly available dataset of total RNA sequencing from Yu *et al* and applied a suite of computational algorithms to create a comprehensive map of circular RNAs in rats. A subset of candidates were further independently validated using experimental approaches. This is by far the most comprehensive circular RNA transcriptome of the rat with respect to tissues, developmental stages and gender using multiple combinations of tools.

## Materials and Methods

### Datasets

A total of 320 datasets were downloaded from the Gene Expression Omnibus, a publicly available database with the accession ID GSE53960. These datasets encompass study by Yu *et al* report the transcriptome from different tissues and developmental stages of the rat. We downloaded ribosomal RNA-depleted RNA-sequencing datasets for different organs including liver, heart, kidney, brain, lung, muscle, spleen, thymus, adrenal gland, uterus and testis. These samples were further categorized as male and female, the age of 2, 6, 21 and 104 weeks and four biological replicates each [GEO: GSE53960]. We have converted Sequence Read Archive (SRA) files to FASTQ format using SRA toolkit for further mapping and analysis. The SRA IDs for the datasets used in the present analysis have been summarised in **Supplementary Table 1**.

### Identification of circular RNAs from Rat samples using CirComPara

We downloaded *Rattus norvegicus* reference genome (version *rn6*)from UCSC genome browser and annotation file from Ensembl(Rosenbloom et al., 2015) database. We used the CirComPara pipeline to identify circular RNAs. CirComPara(Gaffo et al., 2017; Rosenbloom et al., 2015) is an automated pipeline to detect, quantify and annotate circular RNA junctions from RNA-sequencing data and can be used for four different pipelines to identify backsplice junctions. This pipeline also quantifies the expression of linear RNA and their gene expression which can be compared and correlated with circRNAs. CirComPara uses four different detection tools including CIRI, testrealign, CIRCexplorer and find_circ. CirComPara can be used in any pair of pipelines or alignment tools with customized cut-offs for read coverage and filters for alignment scores. We used Segemehl, STAR, Tophat and Bowtie2 aligners, creating a total of 4 combinations - Segemehl_CIRCexplorer2, Star_CIRCexplorer2, tophat_CIRCexplorer2 and Bowtie2_FindCirc. The first step to identify backsplice junctions is to align the RNA-sequencing reads. CirComPara aligns the RNA-sequencing reads over the reference genome using Hisat2 to discard reads aligning over linear transcripts and uses only the discarded reads to identify circ-junctions. After discarding the mapped reads that were possibly mapping with the linear transcriptome, we took the unmapped reads to identify circRNA junctions. We aligned our data using different aligners including hisat, bowtie, segemehl, star and tophat. **Figure 1** shows a schematic representation for identification of circular RNAs.

**Figure 1.**
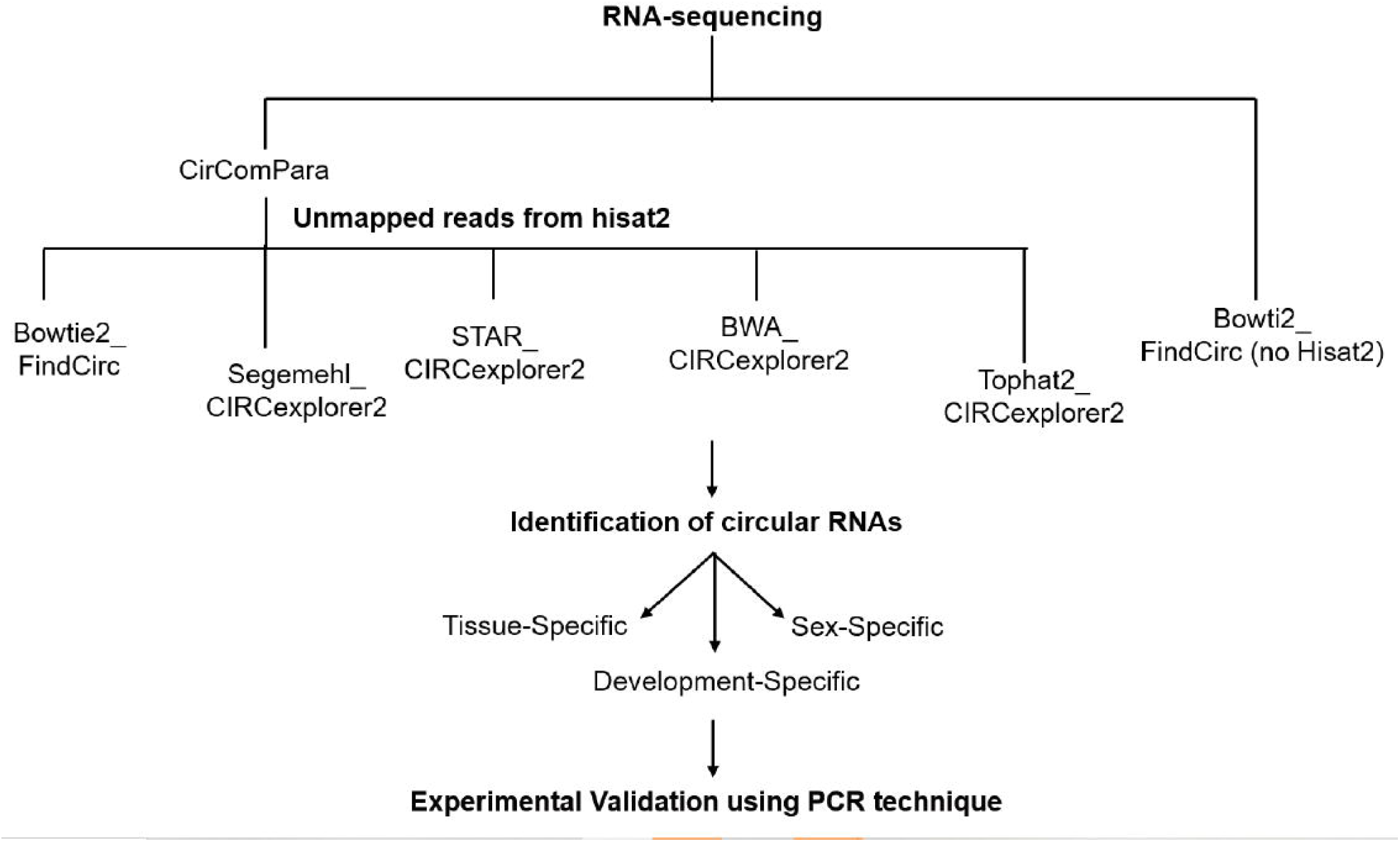
The schematic representation of the pipeline followed for the identification of circular RNAs.

### Identification of circular RNAs from Rat samples using FindCirc

In addition to CirComPara, we have also added find-circ with Bowtie2 as the aligner (the standard pipeline mentioned in Memzak *et al* 2013(Memczak et al., 2013)) as an additional pipeline without using Hisat2 to discard unmapped reads which is default default feature in CirComPara. The datasets downloaded were aligned over the indexed reference genome rn6 using Bowtie2. In this approach, after alignment using Bowtie2, unmapped reads were segmented and mapped from the terminals head and tails to form anchor sequences. We mapped the FASTQ file over the reference genome and discarded the mapped reads. The mapped reads were discarded to avoid any ambiguity with linear transcripts. Next, using the customized scripts from the reference study, we took 20 mers from the head and tail of unmapped reads for unique alignment and extended the read to form an anchor sequence. Certain filters were applied including unique anchor alignment, maximum of 2 mismatches in extended sequences and alignment score above 35, a splice site of GU/AG to avoid false-positives candidates as mentioned in the reference study. The reads that fulfilled the criteria to form putative circular RNAs were saved as output bed files.

### Genome-wide annotation and distribution of circular RNA junctions

We downloaded the annotation GTF file for *Rattus norvegicus* from ensemble (Rnor_6.0.93). The annotation file has 32624 genes, 27780 5’UTR, 22322 3’UTR, 40807 transcripts, 26224 start codon and 25926 stop codon coordinates. We overlapped the circRNA junctions identified from each sample over the annotation file to understand the genome-wide distribution pattern, if any, for circRNA coordinates.

### Tissue-specific, gender-specific and developmental-stage-specific Circular RNAs

We further analyzed the identified circRNA junctions for tissue-specificity. For each tissue, we also analysed the data with respect to developmental stage and gender. We also identified candidates with read coverage >2, >2 to 10<, >10 to 100<, 100-200 and >200 to study the relation of function with their expression level in the tissue.

### Splice-site identification

We extracted the splice site nucleotides for each circRNA junction to analyze any pattern in the splice site of circular RNA and to spot any difference from the linear counterpart. We have studied three conditions as shown in **Supplementary Figure 1**.

### Experimental validation

In order to validate the predicted circRNA junctions from Bowtie2_FindCirc pipeline, we used a polymerase chain reaction (PCR) based approach. We randomly selected candidates based on the significance of genes in the tissue and expression of circular RNA junction that should be expressed in at least 30 samples out of 32 samples for each tissue. In order to validate the circRNA junctions, PCR amplification was performed on cDNAs of corresponding tissues and genomic DNA of the rat using divergent primers. We designed divergent primers of ∼20bp length across the predicted back-spliced junction for different circRNAs in 6 rat tissues (Brain, Heart, Lungs, Liver, Kidney, Thymus). First stranded complementary DNA (cDNA) was prepared from 500ng of RNA from individual tissues using random hexamers and superscript II reverse transcriptase (Invitrogen, USA). PCR amplification was performed using divergent primers designed for a total 24 candidates across 6 tissues using respective cDNAs.

### Expression analysis using quantitative real-time PCR

Expression analysis was performed for selected circular RNA candidates, predicted for their tissue-specific expression patterns. RNA from 6 different tissues (Brain, Heart, Lungs, Liver, Kidney and Thymus) were used to synthesize cDNA as previously described. Circular RNA levels were quantified by quantitative Real-Time Polymerase Chain Reaction (qRT-PCR), using Sybr Green mix (dssTakara, Japan) and detected by Lightcycler LC 480 (Roche). Primer sequences for qRT-PCR have been summarised in **Supplementary Table 2**.

### RNase R treatment to validate circular RNAs

RNA from each of the 6 tissues were treated with RNase R as described previously(Sharma et al., 2019). In order to validate circular RNAs candidates, 5ug of RNA from individual tissues were treated with 15 units of RNAse R (Epicentre, Illumina, USA) for 15 minutes at 37 degrees Celsius. LiCl was used to precipitate the RNAse R treated (RNaseR +) and untreated RNA (RNaseR -). After the purification, cDNA was synthesized from 1ug of the untreated RNA and from the same amount of RNAse R treated RNA using random hexamer and Superscript II reverse transcriptase (Invitrogen, USA). PCR amplification was attempted using divergent primers for selected circular RNA candidates. *Beta-actin* was used as a control, for which convergent primers were used in the experiment. Primer sequences for PCR have been summarised in **Supplementary Table 2**.

## Results

### Summary of RNA-seq data

The RNA-seq dataset used in the study was obtained from a previous publication which sequenced 11 tissues. The data for 320 samples encompassing 11 tissues, 4 developmental stages, both gender (male and female) were retrieved from NCBI SRA. Each of the datasets had 4 replicates each and had an average of 40 million reads each. The samples and the read counts are summarised in **Supplementary Table 1**. The average read count is 41 million reads for which Lvr_21_M_1 had the minimum number of reads, 16038547 and Kdn_104_M_2 had the maximum number of reads that is 82590785. **Figure 2** shows the total number of reads for each sample.

**Figure 2:**
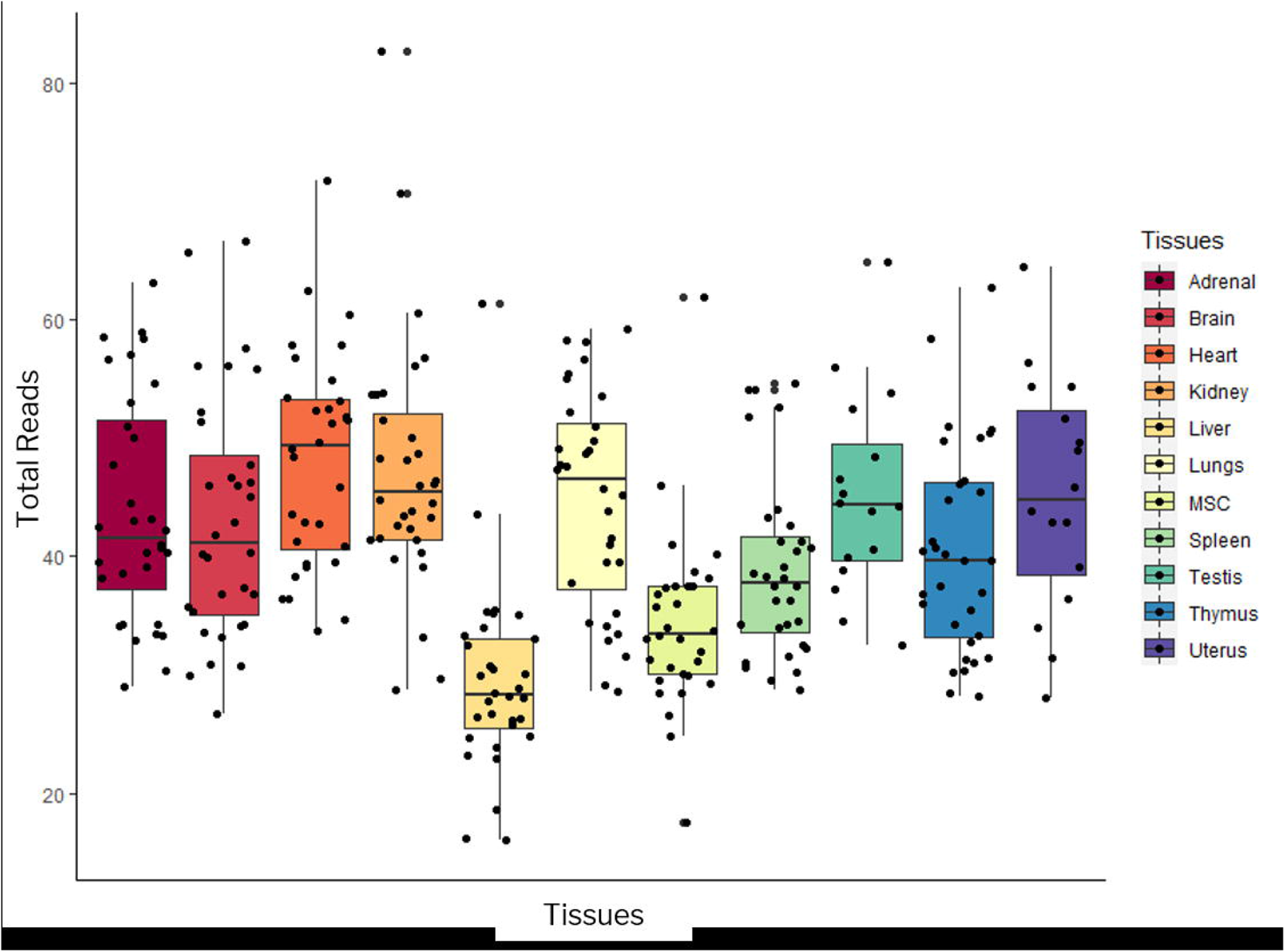
The read numbers in the different tissue sample datasets considered in the analysis.

### Combination of tools used to identify circular RNA junctions

After alignment of RNA-sequencing reads over Hisat2, the average alignment percentage was 87.78%. **Supplementary Table 1** shows the alignment details for each sample using hisat2. The minimum alignment was 68.75% in the liver tissue and maximum alignment was 90.53% in the kidney tissue. The unmapped reads were processed through Segemehl, Tophat, STAR and Bowtie2 aligners to identify backsplice junctions. The alignment details for each aligner have been summarised in **Supplementary Table 1**. The mean alignment in case of Bowtie2_findcirc (noHisat2) was 85%. Similarly, in other cases including Bowtie2, STAR, Segemehl and Tophat, the mean alignment percentage was 66.7%, 38.7%, 73.37% and 61.5% respectively. From the unmapped reads, we observed an alignment of 30-75% from Bowtie2, 42-85% from Segemehl, 20-57% from STAR, 32-75% from Tophat, 15-76% from Bowtie2_FindCirc. **Figure 3** shows the alignment percentage for each sample for each pipeline.

**Figure 3:**
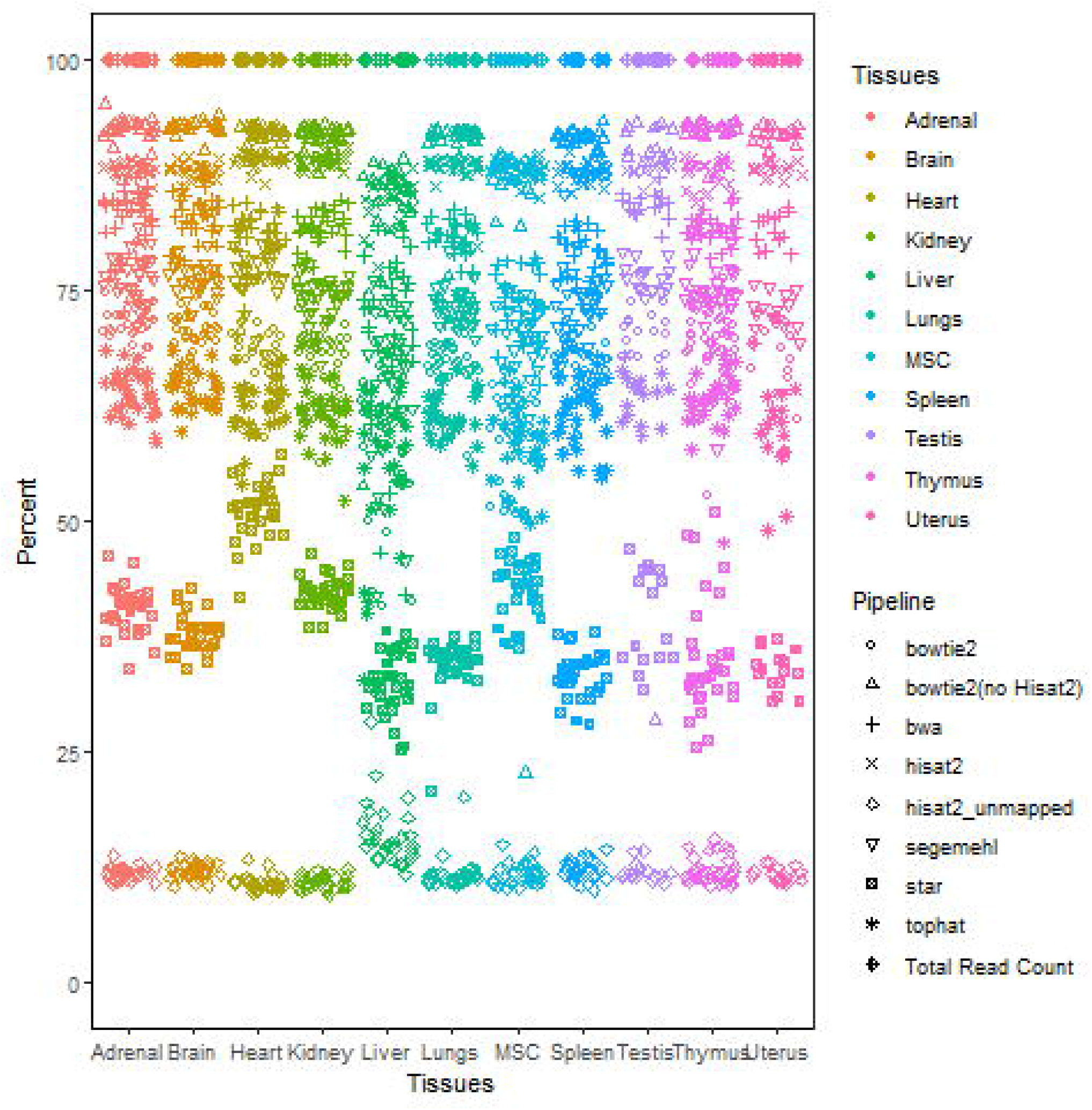
The figure shows the alignment details of the samples in the different combinations of tools.

### Identification of circular RNAs

Our analysis identified a total of 57,022 unique back-splice junctions in the rat genome from Segemehl_CIRCexplorer2, 21,375 from STAR_CIRCexplorer2, 30,943 from Tophat_CIRCexplorer2, 244,511 from Bowtie2_FindCirc and 9,109 from Bowtie2_FindCirc (no Hisat2). Out of the total 5,358,249 unique cirRNA junctions from these 5 combinations, we found 490 were common among all the five combinations **(Supplementary Table 3)**.

### Genome-wide distribution of circ-junctions

Different studies have shown that circular RNAs loci in the genome could map to exons, introns, 5’UTR as well as 3’UTR. We mapped the circular RNAs identified from each combination to the refseq annotations to analyze the overall genome-wide distribution of circ-junctions. For the circular RNAs identified by Bowtie2_FindCirc (no Hisat2), our analysis revealed 1,111 circRNAs mapping (from start to stop) to the 5’UTR, 506 circRNAs to the 3’UTR, 1,361 overlapped with the start codon, 730 overlapped with the stop_codon. In addition to this, we found 86 and 83 circRNAs that had start and stop boundaries upstream 1000 from gene start and downstream 1000bps gene end respectively. In the case of Bowtie2_FindCirc (after Hisat2), we found 3,093 and 3,062 circRNA junctions from upstream and downstream 1,000 base pairs from gene boundaries. Similarly of the circular RNAs identified by the STAR+CIRCexplorer2 approach, 1,994 mapped to the 5’UTR, 369 to the 3’UTR. We could not find intergenic regions in CIRCexplorer2 as it uses existing gene models for annotation by default and we would need to select the “--low-confidence” option. We did not want to increase false positive in our analysis. The numbers overlapping the genome features have been shown in **Supplementary Table 4**.

### Tissue-specific circRNA junctions

Since the dataset encompassed 11 tissues, we further explored the tissue specific circular RNA junctions. Out of the total circular RNAs identified from 11 tissues, the brain was found to have the maximum number of circRNA junctions and the liver was found to have the least number of circRNA junctions. The **Supplementary Table 5** shows the total number of circular RNAs identified from each tissue. Out of the total circRNA-junctions, the unique number of circRNAs specific to each tissue is also mentioned in **Supplementary Table 6**. The data clearly shows tissue enriched and tissue-specific circular RNAs. We have shown the circ-junctions identified from each pipeline specific to tissues in **Supplementary Table 1. Figure 4** summarises the pattern of circRNA-junctions with respect to tissues and pipeline used. While comparing all the combinations, we found that 130 from adrenal, 806 from brain, 208 from heart, 326 from kidney, 308 from lungs, 80 from liver, 111 from MSC, 312 from thymus, 244 from spleen, 299 from testis and 135 from uterus were common in all 5 combinations. From these total circRNAs among 5 pipelines, we found 11 from adrenal, 407 from brain, 37 from heart, 10 from liver, 49 from lungs, 83 from kidney, 61 from thymus, 26 from spleen, 13 from MSC, 97 from testis and 13 from uterus tissue-specific circ-junctions **Supplementary Figure 2**.

**Figure 4:**
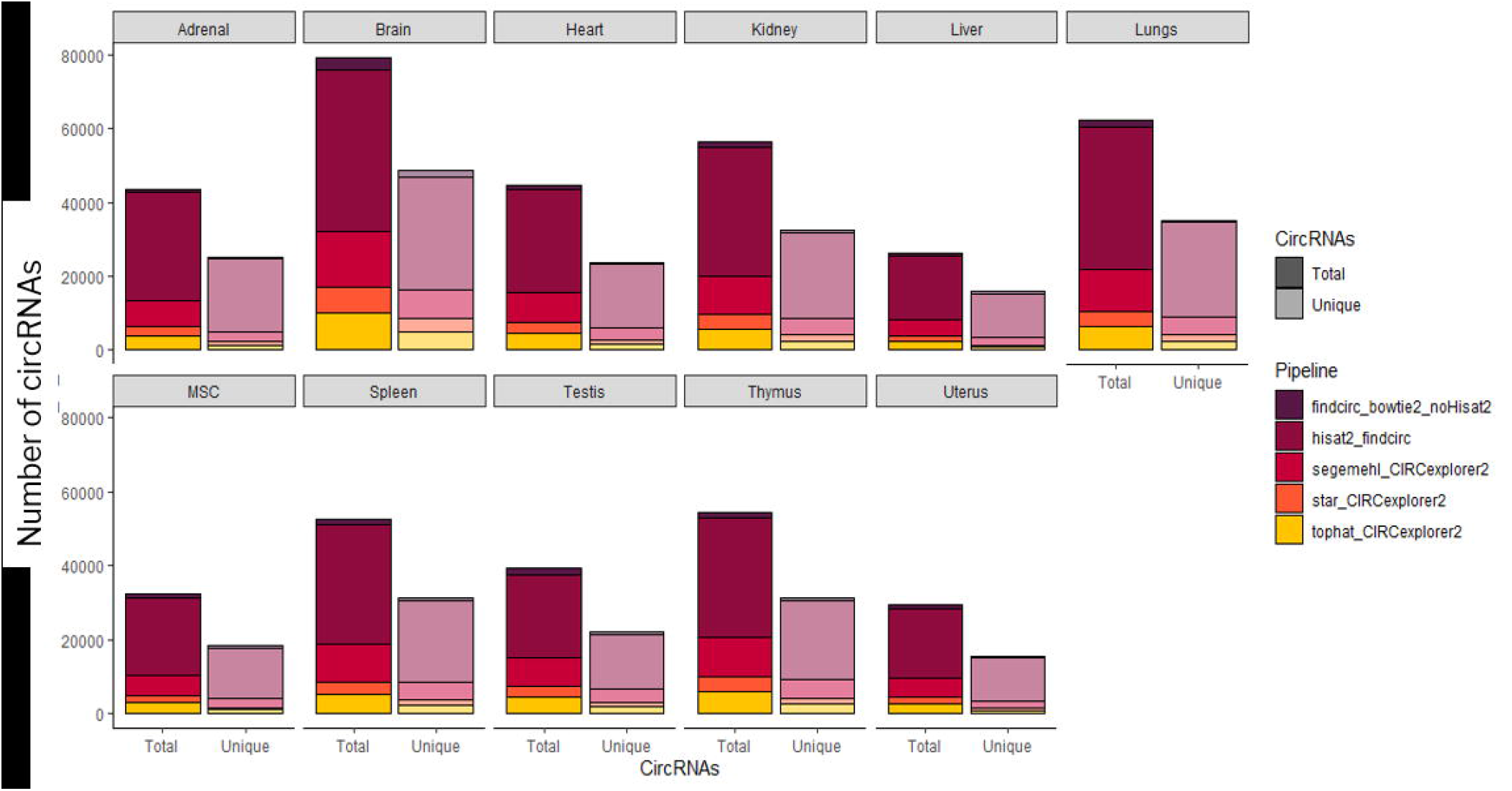
The plot showing the total vs tissue-specific number of circular RNAs for each pipeline. (MSC=Mesenchymal stem cells)

### Development-stage specific circRNA junctions

The RNA-sequencing dataset utilised in this analysis encompassed 4 developmental stages of rats that are 2 weeks-old, 6 weeks-old, 21 weeks-old and 108 weeks-old. Out of the total circ-junctions identified from Segemehl_CIRCexplorer2, we found 10,060 circ-junctions uniquely belonged to 2 weeks-old stage, 10,852 to 6 weeks-old stage, 10,420 to 21 weeks-old and 10,753 to 108 weeks-old stage of the rat. In the case of Bowtie2_FindCirc, we found 1,093 uniquely from weeks 2, 1,241 from weeks 6, 1,580 from weeks 21 and 1,674 from weeks 104. **Supplementary Figure 3** shows the development-stage-specific circRNAs. With respect to tissues, we found a maximum number of 2 weeks-old stage-specific circRNAs in brain tissue and least in uterus tissue. Similarly, in 6 weeks-old stage, we found maximum circRNAs in brain tissue and least in liver tissue followed by 21 weeks-old stage where we found circRNAs in brain tissue and least from liver tissue and 108 weeks-old stage where we found circRNAs in brain tissue and least in mesenchymal cell tissue. **Supplementary Figure 4** shows the tissue-wise development-stage specific circRNAs in rat. Our analysis suggests the maximum number of circular RNAs are identified in 104 weeks stage and least from 2 weeks stage (**Supplementary Table 7-12**). Among all 5 combinations, we found 537 circRNA junctions common at 2 weeks stage, 655 circRNA-junctions at 6 weeks stage, 776 at 21 weeks stage and 833 circ-junctions at 104 weeks stage. Out of these common junctions expressing among all pipeline combinations, we found 102 uniquely expressing at 2 weeks stage, 130 at 6 weeks stage, 193 at 21 weeks stage and 247 at 104 weeks stage and 258 were common among these stages. **Figure 5** shows that the number of circular RNAs identified from each pipeline for each development stage. The venn diagram is shown in **Supplementary Figure 5** that shows the common circular RNAs from all the combinations of tools.

**Figure 5:**
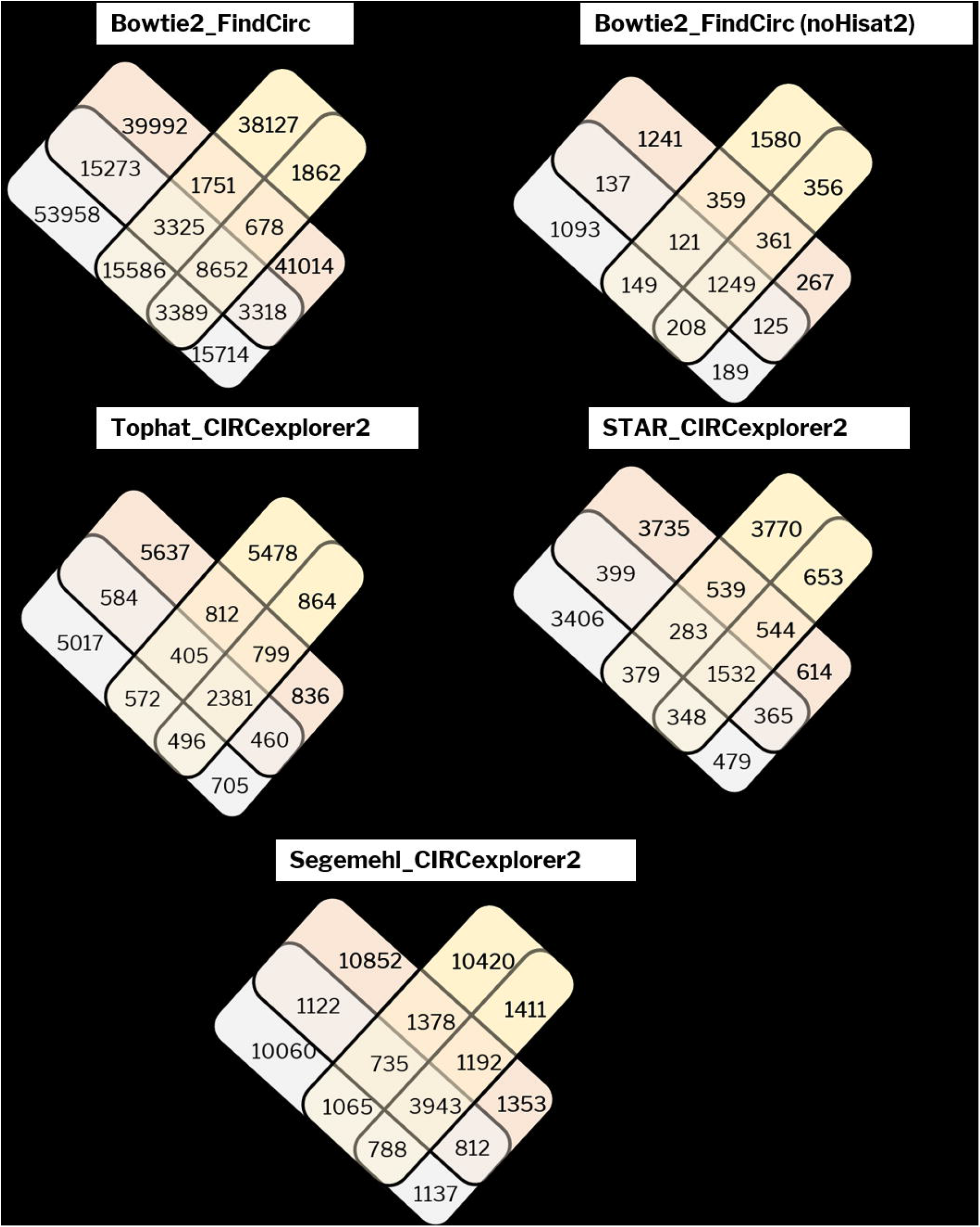
The figure shows a venn diagram comparing each pipeline with different development stages.

### Gender-specific circRNA junctions

We further categorized the circ-junctions based on gender. We identified 5,517 circRNA junctions from female samples of Bowtie2_Findcirc (no Hisat2) and 6,524 circ-junctions from male samples. Out of these 9,109 total unique candidates, 32.18% junctions were common among both genders. We found 46.86% and 55.06% unique circ-junctions from females and male respectively. Similarly, in the case of Bowtie2_FindCirc, we found 79.68% unique circ-junctions from females and 80.23% from male. In the case of Segemehl_CIRCexplorer2, we found 21% unique common junctions. We identified 23.6% common circ-junctions, 40% unique male and 37.72% unique female junctions from Star_CIRCexplorer2. Out of total circRNAs junctions from male, we found ∼89% unique circRNA junctions from male and female both with 11.9% common circ-junctions. **Figure 6** shows the distribution of circular RNAs in both genders. Among the common circ-junctions from 5 combinations, we found 683 circRNA junctions common among all 5 pipelines. We found 59 circ-junctions unique to the male gender and 25 circ-junctions unique to the female gender common in all 5 pipelines (data shown in **Supplementary Table 13-14**)

**Figure 6:**
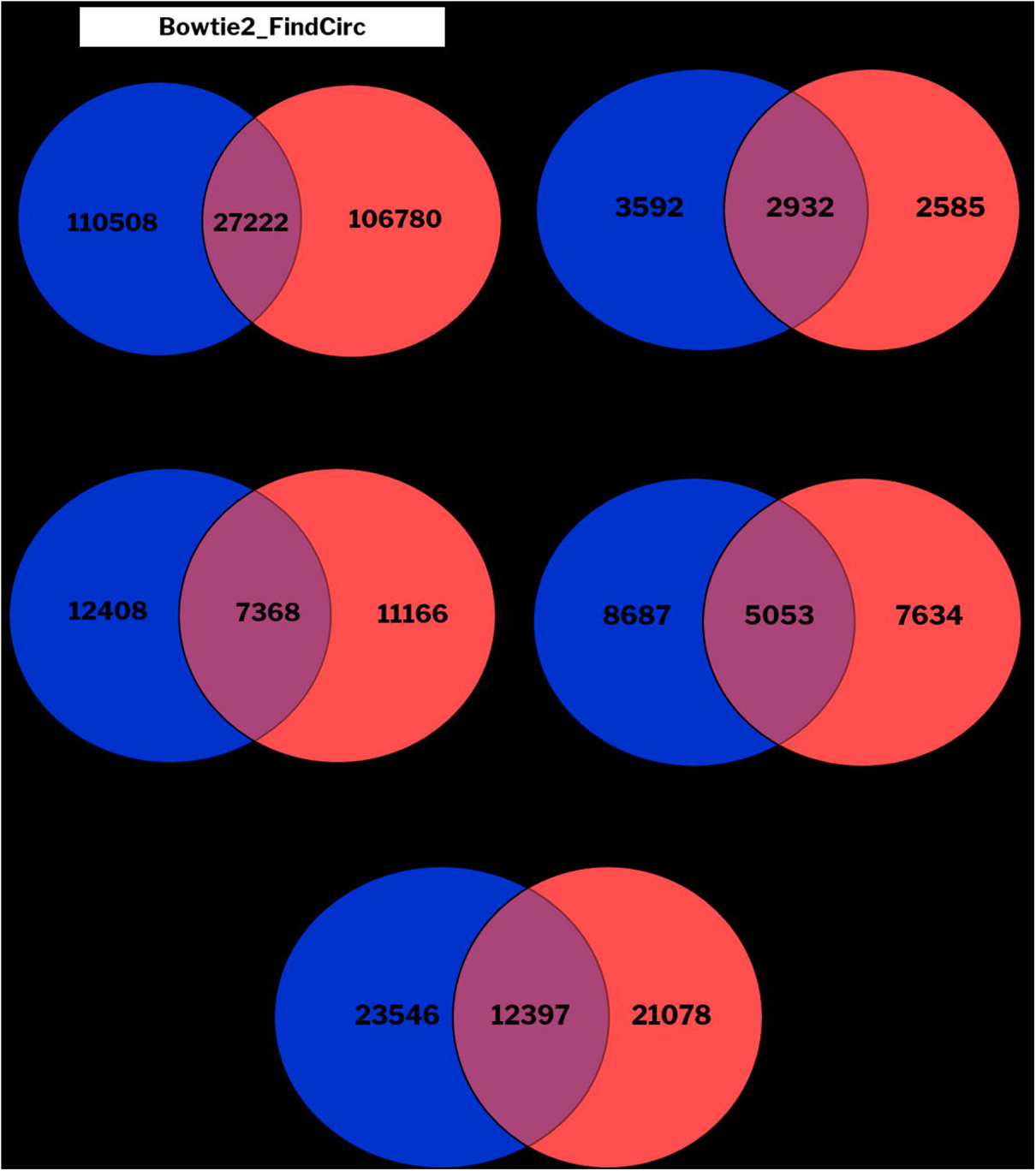
The figure shows the venn diagram for gender-specific circRNAs in each pipeline.* =Bowtie2_findCirc (no Hisat2) includes circular RNAs identified from Memczak et al pipeline without aligning prior with Hisat2.

### Circ-junctions based on read coverage

We have shortlisted circ-junctions with cut-off of read coverage at splice junctions with cut-off >2, >2 and 10<, >10 and 100<, 100-200 and >200 to study the relation of function with their expression level in the tissue. Among 5 different combinations, we found that maximum number of circRNAs with circ-count 26,094 fall with >200 cut-off is from Segemehl_CIRCexplorer2 and minimum circRNAs with 3 from >200 cut-off is from Bowtie2_FindCirc combination. Segemehl_CIRCexplorer2 has a maximum number of reads in the range 100-200 reads. **Supplementary Table 15** shows the read counts from each range for each combination and **Figure 7** shows the overall distribution of reads by coverage cut-off.

**Figure 7:**
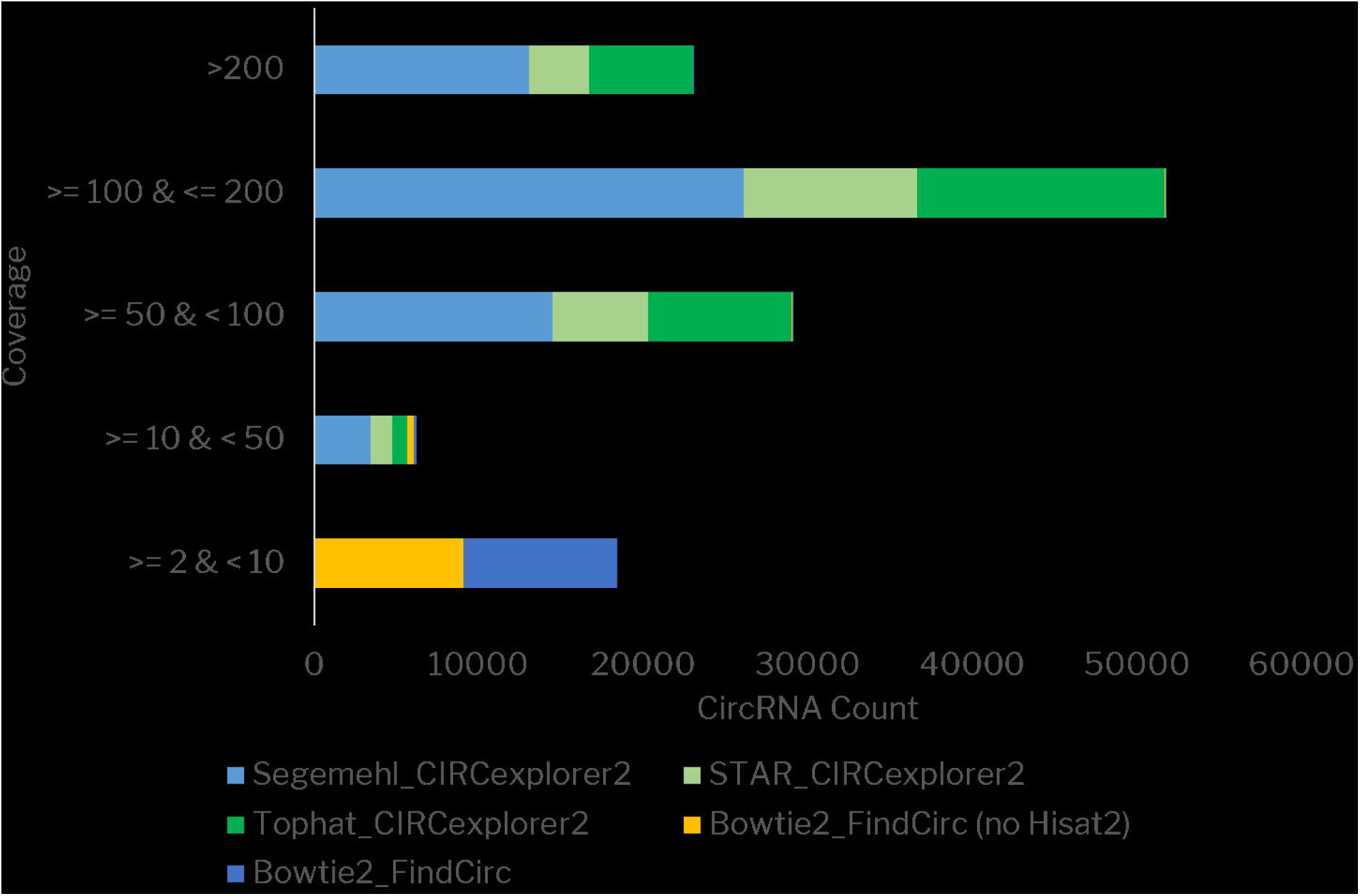
The classification of circular RNAs based on read coverage from each of the pipelines considered.

### Splice-sites involve in the formation of circRNA junction

We identified the splice sites for all the three cases from start and stop sites for each pipeline combination and interestingly we found that splice sites used to identify backsplice junctions for each pipeline were conserved. We found conserved AG and AC splice sites at the start position (case 2) and GG,CC splice sites at stop position. We also found that there is conserved T/C at one site before the splice site (TA and CA at start sites in case 1) and G/A at one site after stop position (CC,GG,GA at stop site in case 3). Similarly, at the stop position, we found conserved C/G right before the splice site (case 1 with CT/GT splice sites at stop site) and A nucleotide one position after the splice site.

### Experimental Validation

We validated 16 circ-junctions out of 24 candidates selected based on literature significance using divergent primers approach with PCR in tissues including brain, thymus, liver, lungs and kidney. The list of selected candidates is given in **Supplementary Table 2** with the primer sequences. Divergent sets of primers of length ∼20 nucleotides were designed overlapping the back-splice junction to obtain the amplicon product of around 200-300 base-pairs. The selected candidates were amplified from cDNA of respective tissues and genomic DNA as control. The genomic DNA was used to negate the possibility that the sequence of circRNA junctions could be a part of DNA itself as there are cases of repetitive exons and transplicing are reported. The results of experimental validation have been summarized in **Figure 8**.

**Figure 8:**
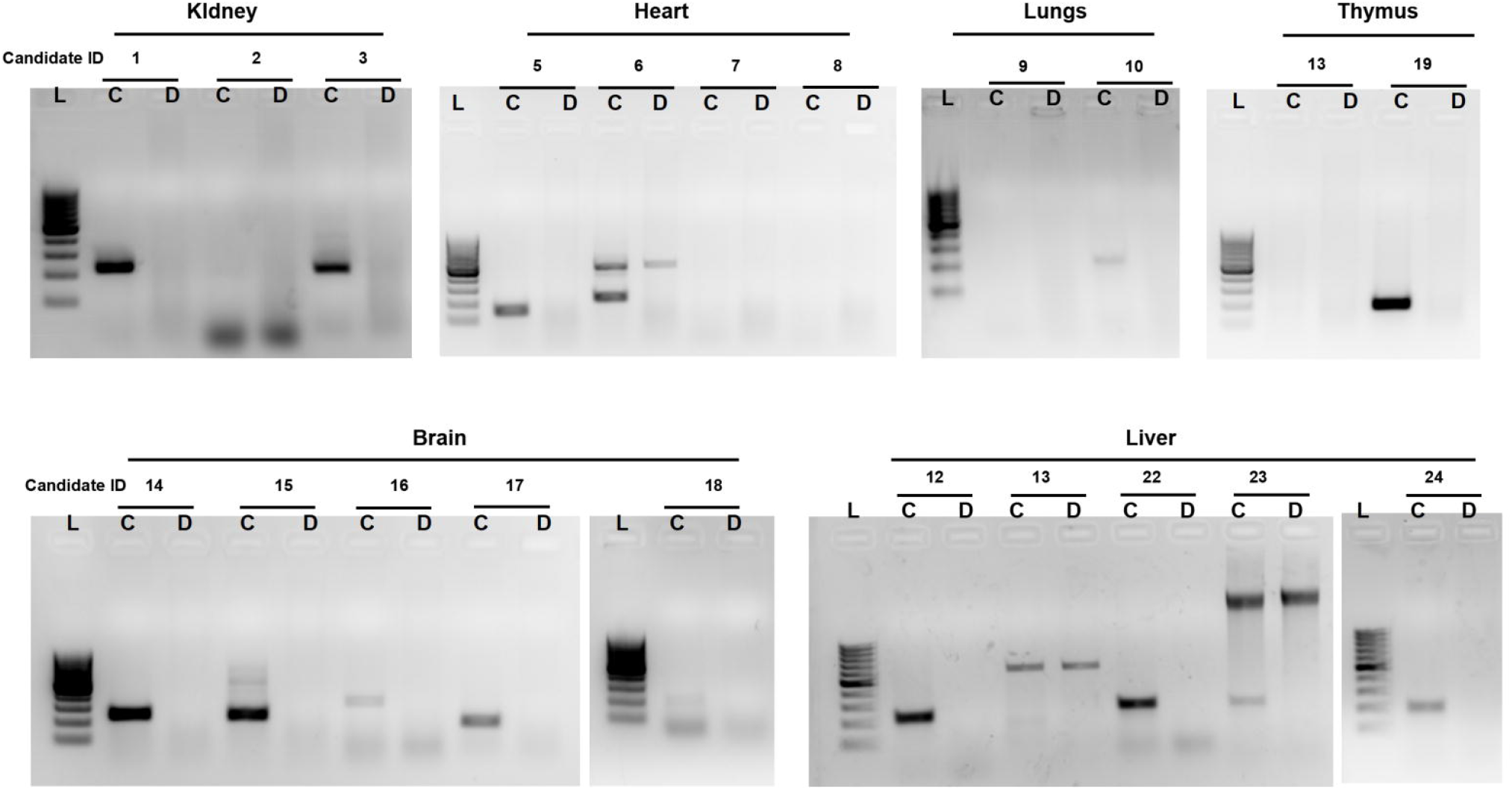
Validation of circular RNAs using RT-PCR for 6 tissues panel. Agarose gel represents the amplification products obtained from cDNA synthesized from different rat tissues and control rat DNA. Total 24 candidates were analyzed in 6 different rat tissues.

### Quantitative expression using qRT-PCR

The experimentally validated circRNA candidates were further analyzed for their quantitative expression analysis using qRT-PCR based approach in these tissues. Interestingly the expression of circRNA from brain specific gene *‘Anks1b’* has shown the tissue-enriched expression in case of brain and the expression of circRNA originating from thymus specific gene *‘Themis’* has shown the tissue-enriched expression in case of thymus. In the case of the liver, *Efemp1* showed tissue-enriched expression. We also performed RT-PCR for 2 circRNAs originating from genes *‘Ubr5’* and *‘Ralgapa1’*. These circRNAs did not show tissue-enriched expression but showed differential expression among different tissues. We have shown the quantitative expression of the circRNAs in **Supplementary Figure 7**. The **Figure** clearly shows the tissue-enriched expression of *circ-Anks1b, circ-Efemp1* and *circ-Themis* and differential expression of *circ-Ubr5* and *circ-Ralgapa1*. The validation of circular RNAs was also done using RNaseR (shown in **Figure 9**). We have also compared the expression with the bioinformatics data from Bowtie2_FindCirc (no Hisat2) read-coverage and the RT-PCR expression was shown to be in coherence with the bioinformatics data. The bioinformatics data for these candidates have been shown in **Supplementary Figure 6a and b**.

**Figure 9:**
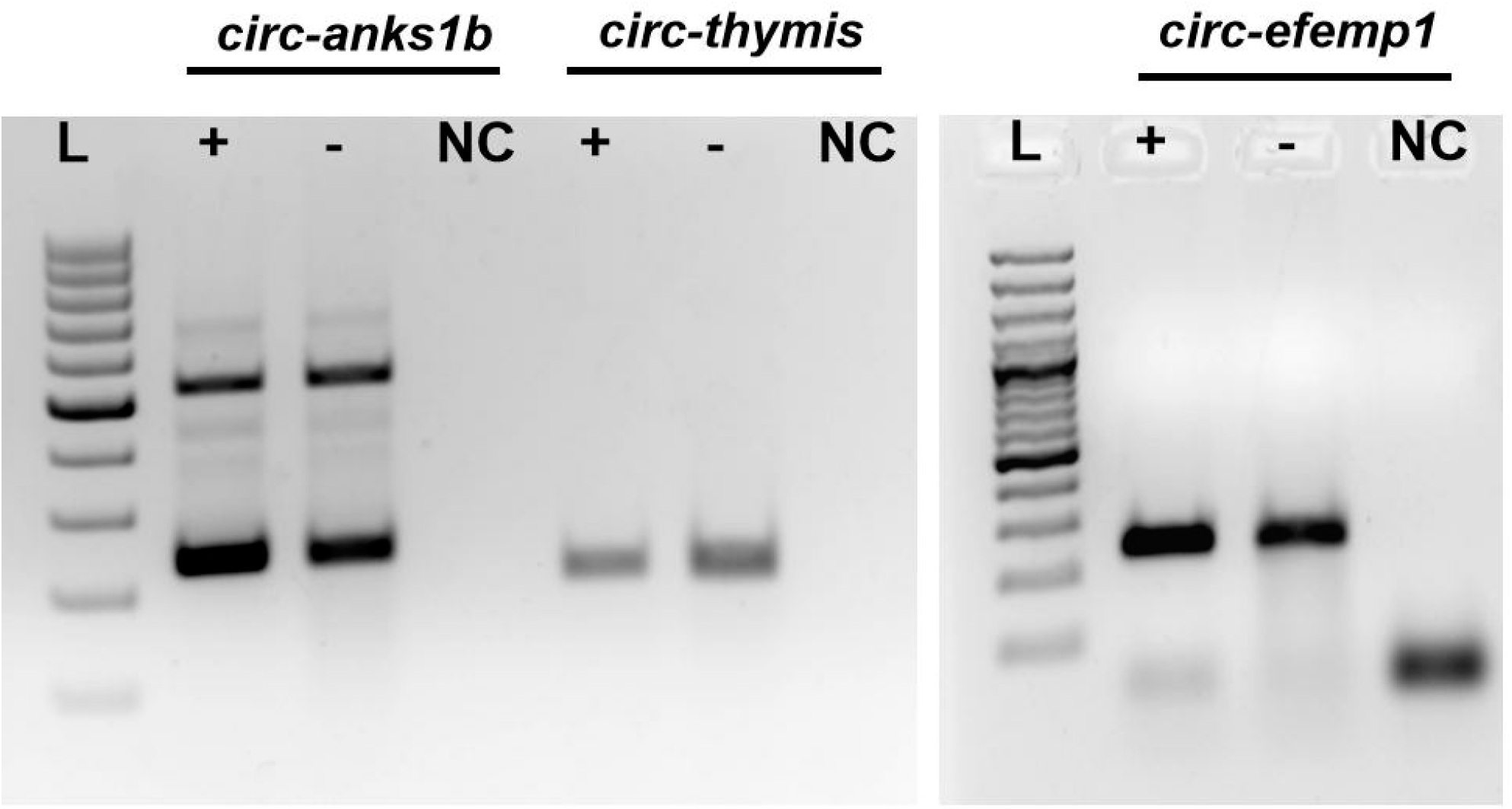
Validation of tissue-enriched circRNAs with RNaseR treatment using PCR based approach Agarose gel represents PCR amplified products in RNaseR treated (+) and RNaseR not treated (-) and no template control (NC).

## Discussion

In this study, we have performed a comprehensive analysis of circular RNAs in rat utilising transcriptome profiling dataset made available by Yu *et al(*Yu et al., 2014). The dataset encompassed 320 samples including 11 tissues, 4 developmental stages and 2 genders each with 4 replicates.

Since no single computational approach is perfect to detect circRNAs at high sensitivity and specificity, we used a combination of tools to identify putative backsplice junctions. Our study focussed on using 5 different pipeline combinations to identify circRNA junctions. These pipelines include tophat_CIRCexplorer2, Star_CIRCexplorer2, Segemehl_CIRCexplorer2, Bowtie2_FindCirc and Bowtie2_FindCirc (no Hisat2). These pipelines use aligners including segemehl, star, Bowtie2 and tophat to identify the backsplice junctions and then different algorithms including CIRCexplorer2, Find_Circ and segemehl to filter backsplice junctions. Our study shows the 1) tissue-specific 2) development stage-specific and 3) gender-specific circRNAs from each pipeline combination.

The RNA-sequencing dataset varying read count from 16 million to 82 million. Liver samples had the lowest read count and kidney, brain and heart were among the samples with the most number of reads with very good quality. CirComPara is one published algorithm that uses the combination of circRNA tools to identify circRNA junctions. The first step, in circompara, is to trim the input RNA-sequencing reads with phred cut-off 30. Our analysis showed that the quality of reads were good enough so no reads were removed. Next step is to align the data over alignment tool to remove the reads aligning the linear transcripts. This step is very critical as it removes the ambiguous reads matching with both linear and backsplice junctions and improves the sensitivity of the circRNA detection by reducing false positives. CirComPara uses hisat2 (hierarchical indexing for spliced alignment of transcripts) as the alignment tool to discard the reads aligning to the linear transcriptome. HISAT2 uses two-way indexing, one is whole genome Ferragina-Manzini (FM) index to anchor each alignment and multiple local FM indexes for rapid extension of the alignments. HISAT2 uses only 4GB memory to run and is one of the fastest alignment tools available for large datasets. After alignment with hisat2, with average alignment of 87%, alignment percentage ranged from 68.75% to 90%. The lowest alignment percentage was for liver samples and kidney and heart were samples that showed very high alignment.

CirComPara uses hisat2 as default aligner to discard the linear mapped reads but in case of Memczak *et al(Memczak et al*., *2013)*, the complete pipeline includes Bowtie2 from the step to remove reads mapping over the linear transcripts to the identification of circRNA junctions using the anchor alignment method. So we added one more variation to the default procedure that is findcirc with Bowtie2 aligner without aligning before with Hisat2. In the latter case, Bowtie2 uses options like --very-sensitivity, --score-min=C,-15,0 to use very stringent conditions to filter out reads mapping contiguously and full length to the genomes. We observed the minimum number of circRNAs from Bowtie2_FindCirc (no Hisat2). The study describing CirComPara also explained that testrealign predicts 86% of the total circRNAs predicted and the difference with other methods is at least ∼5 times more which is due to the reason that testrealign does not perform any post-processing specific for circRNAs. So combining it with other methods can reduce the loose predictions so we have not used testrealign in our pipeline. Hansen *et al* compared 5 prediction tools including circRNA_finder, find_circ, CIRCexplorer, CIRI, and MapSplice to check the sensitivity and number of false positives(Hansen et al., 2016). The data from the study clearly showed the problems with each pipeline. In the case of top candidates, Find_circ performs badly as it is very distracted by the highly expressed linear RNA species. CIRCexplorer is the most reliable to predict top 100 candidates. So we did a further analysis using the individual pipeline as well as the combination of pipelines. Comparing analysis between Bowtie2_FindCirc and Bowtie2_FindCirc (no Hisat2), we found that default features of CirComPara does not use 2 read cut-off and splice-site cutoff of 100kb because of which we have identified very high number of circular RNAs in Bowtie2_FindCirc. Moreover, FindCirc is not dependent on existing models of genome whereas CIRCexplorer2 uses annotated GTF file to annotate circular RNAs. Due to these reasons, Bowtie2_FindCirc has a relatively higher number of circular RNAs than CIRCexplorer2.

While comparing tissue-specific circular RNAs, we found the maximum number of circRNAs were expressed in the brain and least in the liver. The high number of circRNAs in the brain is quite expected from the previous studies and though the least number of circRNAs in the liver could be because of the sample read count also. We found ∼22-30% of unique candidates from each tissue. Interestingly, out of number of circRNAs candidates identified from each pipeline, our analysis showed that very small of circ-junctions were common in all the 5 combinations in each tissue ranging from 80 to 900 (maximum in brain), out of which, 15-400 were unique to each tissue with maximum from brain i.e. 400 which increase the depth of the problem. In comparison to literature and existing databases such as circAtlas(W. Wu et al., 2020) and circfunbase(Meng et al., 2019), we found only ∼7000 circRNAs common. Rest of the circular RNA junctions can be considered novel.

Our comparison of the development stage specific circular RNAs revealed most of the circRNAs from 104 weeks-old stage followed by 2 weeks-old stage. We found an interesting pattern for each tissue over the development stage as the number of circRNAs increases over the age in the brain which is also mentioned by Zhou *et al(T. Zhou et al*., *2018)*. Adrenal circRNA count increases from 2 to 21 weeks-old stage then decreases with aging. Heart and thymus have the most number of circRNAs at puberty i.e. 6 weeks-old stage followed by 2 weeks-old stage then reduce with adult stage and aging. Lungs circRNAs are highest at immature stage and aged stage. Interestingly the circRNAs in mesenchymal stem cells are reduced with aging and in Testis, circRNAs are highest in the adult stage and uterus circRNAs increase with age (highest at 104 weeks-old stage). So clearly there is pattern and tissue-specific behaviour in circular RNAs. We also identified gender-specific circRNAs and found only 59 circRNAs that were common in all the circRNAs pipeline combinations and yet unique to the male gender. Similarly, we found 25 circRNAs specific to females that were common in all 5 combinations of tools.

Next, we were interested in observing how every pipeline differs on the level of read coverage and we found contrasting patterns between CIRCexplorer2 and FindCirc as CIRCexplorer2 had most of the circRNAs fall in the range of <200 reads whereas FindCirc pipeline had most circular RNAs in range of 2 −10 reads. Studies have shown that circular RNAs are originating from exons, introns, 5’UTR and 3’ UTRs(Santer et al., 2019). We were also interested to observe the pattern in rat and found that there are many circular RNAs originating from only 5’UTR and from 3’UTRs. This explains the regulatory role of circRNAs in transcription. Even though we did not use arguments to identify novel circular RNAs from CIRCexplorer2, we identified many circular RNAs originating from intergenic regions from FindCirc and CIRI.

We observed differential expression of circular RNAs among tissues. *circ-themis* is one such gene that shows tissue-specific expression in thymus. Host *themis* gene plays a regulatory role in T-cell positive and negative selection during thymocyte development(Chabod et al., 2012; Iwata et al., 2010). Another tissue specific circular RNA is *circ-anks1b* that is predominantly found in the brain and we also found expression of *Anks1b* in brain tissue only. The Anks1b gene protein interacts with amyloid beta protein precursor and is involved in brain development(Tindi et al., 2015). Role of *Efemp1* has also been reported in liver cancer and methylation of the promoter cause decrease in expression of Efemp1 in hepatocellular carcinoma (HCC)(Y. Chen et al., 2018; Dou et al., 2016; Hu et al., 2019). The gene is also shown to be involved in pathogenesis of Alzheimer’s disease. Genes *Ubr5* and *Ralgapa1* are involved in signalling pathways and we have observed circular RNAs originating from these genes expressing in multiple tissues but differentially expressed.

## Conclusions

We have created a comprehensive map of rat circular RNA transcriptome from 320 samples from 11 tissues, 4 developmental stages and 2 gender. We have also validated a few circular RNAs candidates. Out of these, 3 circular RNAs showed tissue-enriched patterns and 2 are differentially expressed.

## Acknowledgements

DS would like to thank Intel India fellowship for providing fellowship. PS would like to acknowledge CSIR for his fellowship. The authors would like to acknowledge Samatha Mathew for her help in performing experiments.

